# Determining the replication kinetics and cellular tropism of the ruminant-associated Influenza D virus on primary human airway epithelial cells

**DOI:** 10.1101/395517

**Authors:** Melle Holwerda, Laura Laloli, Isabel Stürmer, Jasmine Portmann, Hanspeter Stalder, Ronald Dijkman

**Author notes:** These authors contributed equally to this article (Laura Laloli, Isabel Stürmer). Correspondence: Ronald Dijkman, Institute of Virology and Immunology, Department of infectious Diseases and Pathobiology, Vetsuisse Faculty, University of Bern, Länggassstrasse 122, 3012 Bern, Switzerland.

## Abstract

Influenza viruses are notorious pathogens that frequently cross the species barrier with often severe consequences for both animal and human health. In 2011, a novel member of the *Orthomyxoviridae* family, Influenza D virus (IDV), was identified in the respiratory tract of diseased swine. Epidemiological surveys revealed that IDV is distributed worldwide among livestock and that IDV-directed antibodies are detected in humans with occupational exposure to livestock. To identify the transmission capability of IDV to humans, we determined the viral replication kinetics and cell tropism using an *in vitro* respiratory epithelium model of humans. The inoculation of IDV revealed efficient replication kinetics and apical progeny virus release at different body temperatures. Intriguingly, the replication characteristics of IDV revealed many similarities to the human-associated Influenza C virus, including the cell tropism preference for ciliated cells. Collectively, these results might indicate why IDV-directed antibodies are detected among humans with occupational exposure to livestock.

**One-sentence summary of the conclusion:** We show that the ruminant-associated Influenza D virus has direct transmission capability to humans.

## Introduction

After the initial discovery of Influenza D virus (IDV) in 2011, among swine with Influenza-like symptoms, knowledge about this new genus in the family of *Orthomyxoviridae* is increasing (1, 2). Epidemiological studies have shown that the virus has a worldwide distribution, whereby at least two distinct genetic lineages are cocirculating and reassorting (3–10). Because of the high seroprevalence, cattle is the proposed natural reservoir of IDV, in which IDV causes mild respiratory disease symptoms (11). In addition to cattle, IDV-specific antibodies have been detected in swine, feral swine, equine, ovine, caprine and camelid species, suggesting a broad host tropism for IDV (3, 4, 9, 12, 13). However, the most striking observation is the detection of IDV-directed antibodies among humans with occupational exposure to livestock (14).

There are several indicators that IDV has a zoonotic potential. For instance, the utilization of the 9-*O*-acetyl-*N*-acetylneuraminic acid as a receptor determinant, that allows the hemaglutinin esterase fusion (HEF) glycoprotein of IDV to bind the luminal surface of the human respiratory epithelium (1). Interestingly, the utilization of this receptor is also described for the closely related, human associated Influenza C virus (ICV) (15, 16). Furthermore, the detection of IDV-directed antibodies among humans with occupational exposure to livestock and the molecular detection of IDV in a nasopharyngeal wash of a field worker with close contact to livestock indicates that cross species transmission occurs (14, 17). However, thus far, there is no indication of wide spread prevalence among the general population although the virus has been detected during molecular surveillance of aerosols collected at an international airport (18, 19). Therefore, it remains unclear whether IDV can indeed infect cells within the human respiratory tract and thus whether it has a zoonotic potential.

The respiratory epithelium is the main entry port for respiratory pathogens and is therefore and important first barrier for intruding viruses. For more than 15 years, the human well-differentiated airway epithelial cell (hAECs) culture model has been applied as an *in vitro* surrogate model of the *in vivo* respiratory epithelium to investigate a wide range of emerging and zoonotic respiratory viruses on their capability of direct transmission to humans (20–24). The aim of this study is to investigate the transmission capability of IDV to humans by inoculating hAEC cultures with the ruminant-associated IDV. In addition, we sequentially passaged IDV further on naïve hAEC to determine whether infectious progeny virus is produced. This revealed that IDV is able to efficiently replicate in hAEC cultures and can be subsequently passaged. Moreover, due to the similarity of IDV with the human associated IDV, we compared their viral kinetics and cell tropism. This showed that both viruses have similar replication kinetics and share a cell tropism preference towards ciliated cells. These results emphasize that there is no fundamental restriction of IDV replication within the human respiratory epithelium. Therefore, these findings might explain why IDV-specific antibodies can be detected in humans with occupational exposure to livestock.

## Results

As a first step to address the transmission capability of IDV to humans we inoculated the prototypic D/Bovine/Oklahoma/660/2013 strain on hAECs of three biological donors. Viral progeny release was monitored by collecting washes with 24-hour intervals for a duration of 72 hours. To analyse temperature dependent effects, incubation of the cultures was performed at temperatures that correspond with those of the human upper and lower respiratory tract, 33°C and 37°C respectively. The release of viral progeny from the apical washes was analysed by quantitative real-time reverse transcription PCR for viral transcripts and virus titration for infectious virus. The first viral transcripts were detected at 24 hours post-infection (hpi) among all donors, independently of the incubation temperature **(Figure 1A and B)**. However, some temperature dependent differences were observed when the infectivity of the progeny virus was analysed. When incubated at 33°C, viral titres were detected for every donor at 48 and 72 hpi, but only one donor show a viral titre at 24 hpi **(Figure 1C)**. In contrast, we observed viral titres as early as 24 hpi for IDV infection at 37°C for every donor that increased over time **(Figure 1D)**. These results indicate that IDV kinetics seems to be more efficient at ambient temperatures corresponding to the human lower respiratory tract. This, most likely, reflect the necessity for IDV to replicate at the body temperature of cattle, which is between 37 - 39°C.

**Figure 1:**
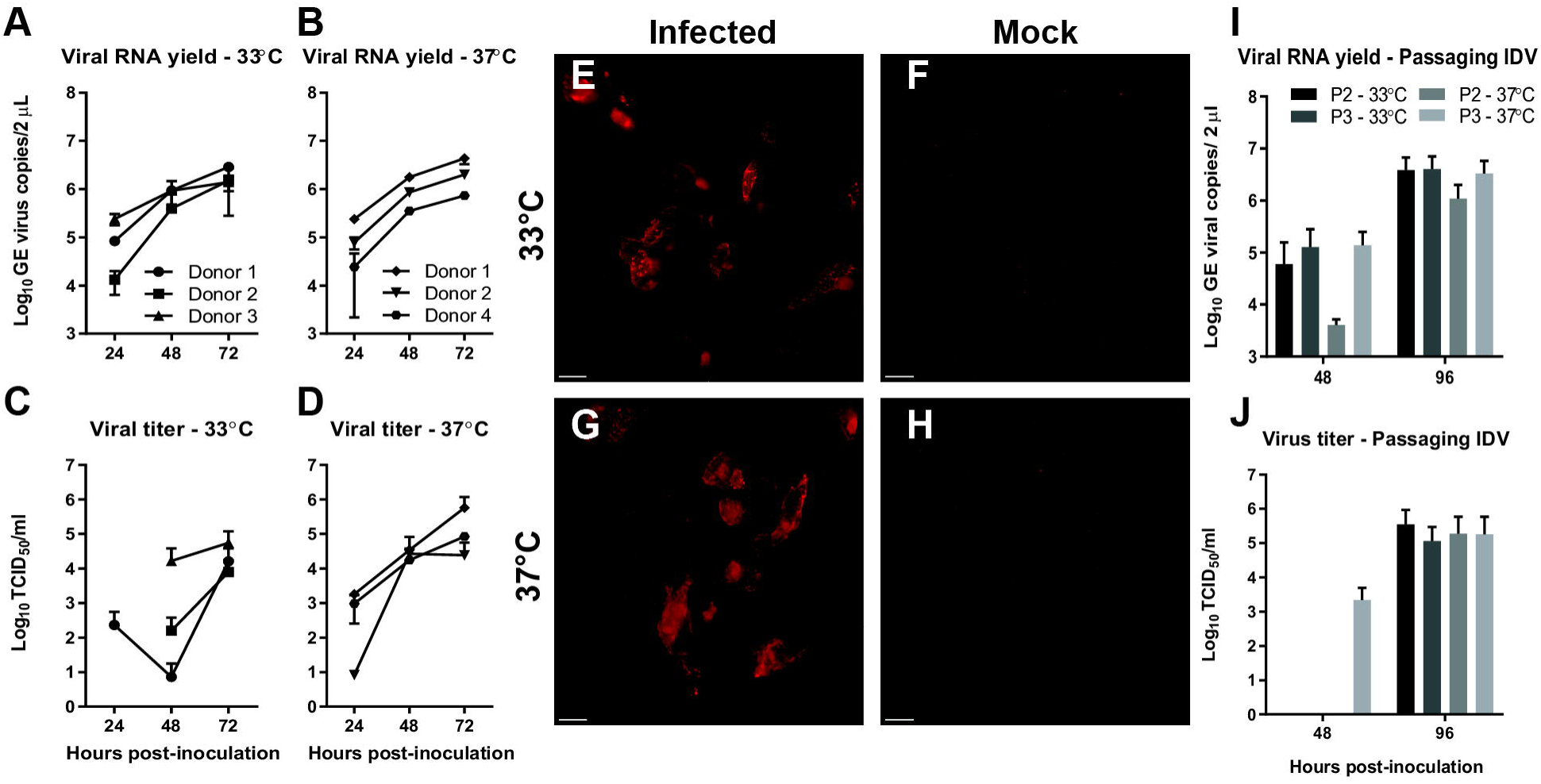
Efficient replication of Influenza D virus (IDV) in hAEC cultures. Human airway epithelial cell cultures were inoculated with 10.000 TCID_50_ of IDV and incubated at either 33°C or 37°C. The monitored viral RNA yield is given as genomic equivalents (GE) per 2 μL of isolated RNA (y-axis) at indicated hours post-inoculation (x-axis) for 33°C **(A)** and 37°C **(B)**. Whereas the viral titer is given as TCID_50_/mL (y-axis) for 33°C **(C)** and 37°C **(D)** at indicated hours post-inoculation (x-axis). These results are displayed as means and SD from duplicates from three independent donors. Human airway epithelial cell cultures were formalin-fixed and immunostained with a custom generated antibody against the Nucleoprotein (NP) of Influenza D virus to detect viral antigen. A representative image from one of the three independent donors is shown for IDV infection at 33°C and 37°C **(E&G)** as well as their respective controls **(F&H)**. Magnification 60x, the scale bar represents 10 micrometer. To assess if IDV viral progeny is infectious, hAEC cultures were inoculated with tenfold-diluted apical wash and sequentially propagated upon new hAEC cultures. The monitored viral RNA yield is given as genomic equivalents (GE) per 2 μL of isolated RNA (y-axis) at indicated hours post-inoculation (x-axis) for each of the conditions **(I)**. Whereas the viral titer is given as TCID_50_/mL (y-axis) for each condition, at indicated hours post-inoculation (x-axis) **(J)**. The results are displayed as means and SD from duplicates from three independent donors.

After having demonstrated that IDV is able to replicate in hAEC cultures from different donors at both 33°C and 37°C, we wanted to corraborate these results via immunofluorescence analysis. However, commercial antibodies against IDV are currently unavailable, so we ordered a custom generated antibody directed against the nucleoprotein (NP) of the prototypic D/Bovine/Oklahoma/660/2013 strain. Microscopic analysis of IDV-infected hAEC cultures revealed clusters of NP-positive cells at both 33°C and 37°C, whereas no fluorescence signal was observed in the control hAEC cultures **(Figure 1E-H)**. The majority of the fluorescence signal from the NP-positive cells has a cytoplasmic distribution pattern, but some of those also appeared to have a nuclear staining pattern. These findings suggest that, like other *Orthomyxoviruses*, the NP of IDV is actively translocated to the nucleus during viral replication (25, 26). Combined, these results demonstrate that IDV is able to efficiently replicate in hAEC cultures from different donors at temperatures corresponding to both the upper and lower respiratory tract of humans.

To analyse if IDV progeny virus is able to infect naïve hAECs of a new donor, we sequentially passaged a 10-fold dilution of the previous obtained 72 hpi apical wash from our three donors on a naïve donor (P2). That was further subpassaged at 48 hpi upon hAEC cultures of the same donor (P3). Like before, we performed the experiment at both 33°C and 37°C to assess whether there are temperature dependent effects. We monitored the production of viral progeny at 48 and 96 hpi for each of the sub-passaging experiment. In the first passage, viral RNA was detected at 48 hpi and increased with one order of magnitude at 96 hpi **(Figure 1I)**. However, we observed that the viral yield at 37°C was approximately one order lower in the first round compared to the viral yield at 33°C, while at 96 hpi this difference slightly reduced. Interestingly, no difference between the different incubation temperatures was observed in the second passaging experiment **(Figure 1J)**. Also, no pronounced differences were observed in the viral titres between the different temperatures or passage numbers at 96 hpi (**Figure 1J)**. However, at 48 hpi, we only detect infectious virus in the apical wash from the last passaging experiment that was performed at 37°C (P3; **Figure 1J**). These results show that the viral progeny from the initial experiments on hAEC cultures is infectious and that IDV can be sequentially passaged on hAEC cultures from different donors at both 33°C and 37°C.

Due to the structural similarity between the HEF of IDV and ICV, and the fact that ICV is a well-known common cold virus that is able to cause a mild upper respiratory tract infection in humans, we wondered how IDV replication efficiency relates to ICV in our hAEC cultures (27, 28). To address this question, we inoculated hAEC cultures with equal amounts of hemagglutination units for ICV (C/Johannesburg/1/66) and IDV and incubated the cultures at 33°C. The viral replication kinetics were monitored as before, by collecting apical washes every 24 hours for a duration of 72 hours. We observed similar replication kinetics for both viruses, although the viral RNA yield for ICV was higher compared to IDV **(Figure 2A and B)**. The replication kinetics of the IDV-infected hAEC cultures were similar compared to the previous experiment at 33°C **(Figure 1A)**. This shows that the replication kinetics for IDV in hAEC cultures is robust and independent from the donor. However, more importantly, we showed that the replication kinetics of IDV are almost identical to that of ICV.

**Figure 2:**
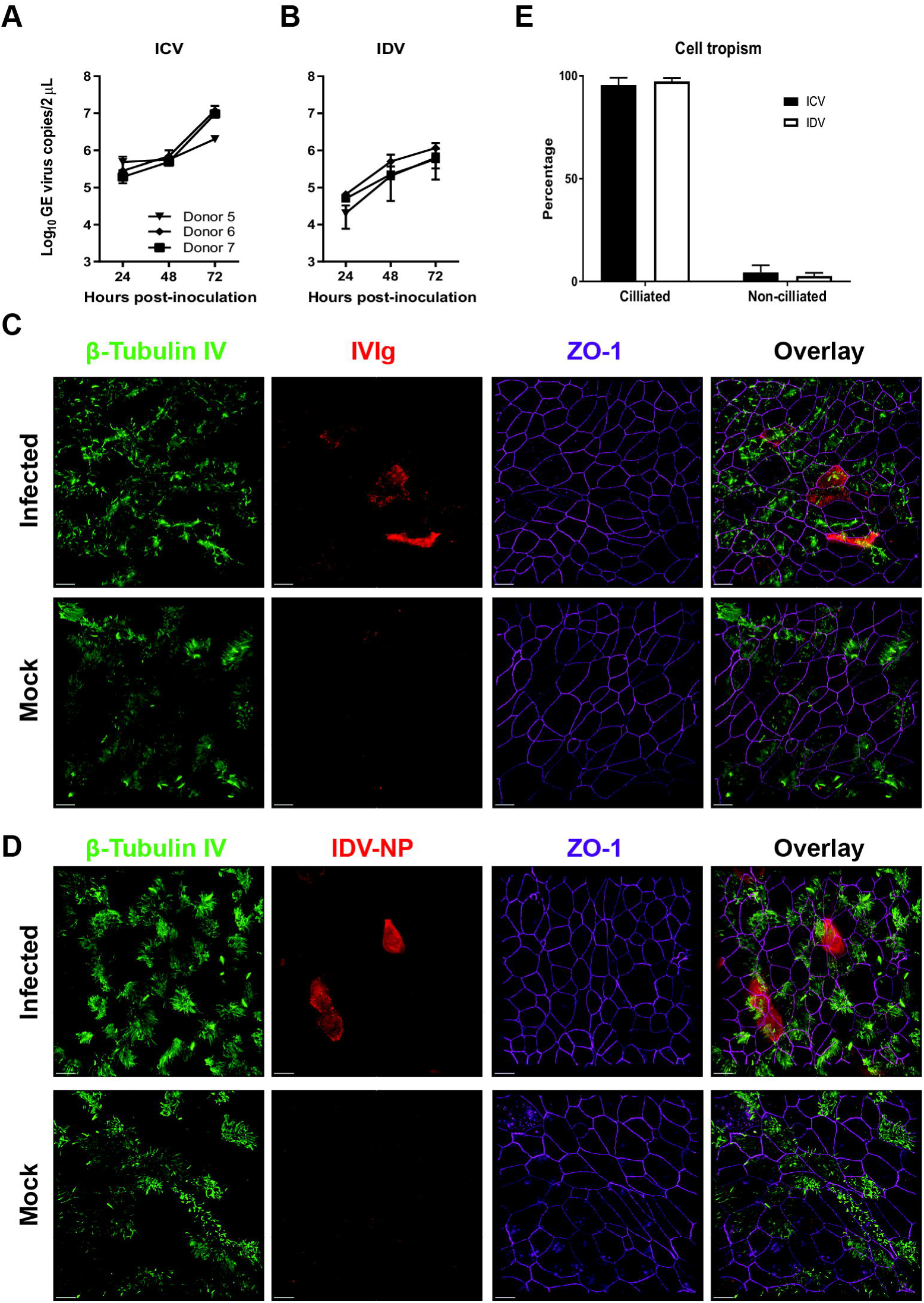
Comparison of ICV and IDV infection in hAEC cultures. Human airway epithelial cell cultures were inoculated with 32 Hemagglutination assay units of ICV or IDV and incubated at 33°C. The monitored viral RNA yield is given as genomic equivalents (GE) per 2 μL of isolated RNA (y-axis) at indicated hours post-inoculation (x-axis) for ICV **(A)** and IDV **(B)**. The results are displayed as means and SD from duplicates from three independent donors. Formalin-fixed ICV and IDV infected hAEC cultures and their respective controls were immunostained with antibodies to visualize the cilia (β-tubulin IV, green), tight junction borders (ZO-1, purple). Whereas virus-infected cells (red) were visualized with either a custom generated IDV NP-antibody or intravenous immunoglobulins (IVIg) for ICV **(C&D)**. Magnification 60x, the scale bar represents 10 micrometer. The cell tropism of ICV (Black bars) and IDV (white bars) was quantified by calculating the percentage of viral antigen-positive signal co-localization with either ciliated or non-ciliated cells **(E)**. The mean percentage and SEM from ten random fields from three independent donors are displayed.

In addition to the replication kinetics, we wanted to determine the respective cell tropism for ICV and IDV, as both viruses utilize 9-*O*-acetyl-*N*-acetylneuraminic acids as receptor determinant (1, 15). We therefore formalin-fixed the infected hAEC cultures to determine the cell tropism for both viruses via immunostaining. To discriminate between the ciliated and non-ciliated cell types, we used well-defined antibodies to visualize the cilia (β-tubulin IV), tight junction borders (Zonula occludens-1, ZO-1) and the nucleus (DAPI). We used our IDV-NP-antibody for detection of IDV infected cells, whereas for ICV we used the commercially available pooled human intravenous immunoglobulins (IVIg). We used the IVIg since most people have encountered one or multiple ICV infections during their life (28, 29). By overlaying the different cellular marker stains with that of the virus antigen, we observed that for both ICV and IDV the virus-antigen signal overlaps with that of the ciliated cell marker **(Figure 2C and 2D)**.

To accurately define the cell tropism, we counted all cell types among ten random fields per donor, with the criteria of at least having one virus-positive cell. For the IDV-infected hAEC cultures, we counted a total of 2273 cells from which 94 were NP-positive, while for ICV a total of 2526 cells and 84 ICV antigen-positive cells was observed. The majority of antigen-positive cells for both IDV and ICV overlapped with the ciliated cell marker with an overall percentage of 97.3 and 95.5, respectively **(Figure 2E)**. This is in line with our initial observation and shows that both IDV and ICV have a predominant preference for ciliated cells. In addition to the cellular tropism, we also calculated the overall infection rate for IDV and ICV, which is 4.1 and 3.3 percent, respectively **(Supplementary table 1)**. These infection rates are in accordance with the previous observed replication kinetics **(Figure 2A and B)**.

Collectively, our results demonstrate that IDV and ICV have similar kinetics in hAEC cultures and share a cell tropism preference towards ciliated cells. Most importantly, these results show that there is no intrinsic impairment of IDV propagation within the human respiratory epithelium.

## Discussion

In this study, we demonstrate that IDV replicates efficiently in an *in vitro* surrogate model of the *in vivo* respiratory epithelium at ambient temperatures that correspond to the human upper and lower respiratory tract. We also demonstrate that IDV viral progeny is replication competent as it could be efficiently sequentially propagated onto hAEC cultures from different donors at both 33°C and 37°C. Intriguingly, the replication characteristics of the ruminant-associated IDV revealed many similarities to the human-associated ICV, including the cellular tropism for ciliated cells. These results show that there is no intrinsic impairment of IDV propagation within the human respiratory epithelium.

For successful inter-species transmission, a virus needs to overcome several barriers before it can efficiently replicate in the new host species (30). These barriers can be classified into three major groups; (i) viral entry through availability of the cellular receptor and proteases, (ii) viral replication and subversion of the host innate immune system followed by (iii) viral egress and release of infectious progeny virus. Our results clearly demonstrate that IDV fulfils most of these criteria for humans, as there is no fundamental restriction for viral replication and sequential propagation of IDV within hAEC cultures from different donors. However, we cannot assess whether IDV can be transmitted between humans with our model. Nonetheless, it has been demonstrated that IDV can be transmitted between both guinea pigs and ferrets, of which the latter is a surrogate model for assessing the transmission potential of emerging Influenza A viruses among humans (31–35). This knowledge in combination with the detection of IDV in aerosols collected at an international airport, and the limited epidemiological data of IDV prevalence among humans, warrants the need for increased surveillance of IDV among humans (18, 19).

At least two distinct genetic lineages are described for IDV, which have over 96% homology, from which the HEF glycoprotein (96.7 to 99.0% homology) is the most divergent of all 7 segments (2, 10). Because cattle are proposed as the main reservoir, we first selected to use only the prototypic D/Bovine/Oklahoma/660/2013 strain, and therefore at that time did not include the prototypic D/Swine/Oklamhoma/1334/2011 strain as a representative of the other lineage. However, due to strict national import regulations for animal pathogens, we currently cannot assess whether both circulating lineages of IDV exhibit similar characteristics in human respiratory epithelium. Although, it is worth mentioning that IDV has been detected in a nasopharyngeal wash of a field worker with close contact to swine (17). Suggesting that both lineages might exhibit similar characteristics in the human airway epithelium.

Both IDV and ICV utilize the 9-*O*-acetyl-*N*-acetylneuraminic acid as their receptor determinant for host cell entry (15, 27). We have shown that both viruses have a predominant affinity towards ciliated cells, suggesting that the distribution of this type of sialic acid is limited to ciliated cells within our *in vitro* model. This tropism is similar to what we previously observed for the human Coronavirus OC43, from which it has been reported to also utilize the 9-*O*-acetyl-*N*-acetylneuraminic acid as receptor determinant (36, 37). Nonetheless, whether this cell tropism for both IDV and ICV corresponds to that of the *in vivo* airway epithelium remains to be determined. Although, we previously have demonstrated that the hAEC cultures recapitulates many characteristics of the *in vivo* airway epithelium, including receptor distribution (36, 38).

In summary, we demonstrate that IDV replicates efficiently in an *in vitro* surrogate model of the *in vivo* respiratory epithelium. This shows that there is no intrinsic impairment of IDV propagation within the human respiratory epithelium and might explain why IDV-directed antibodies are detected among humans with occupational exposure to livestock.

## Material and methods

### Cell culture

The Madin-Darby Bovine Kidney (MDBK) cells were maintained in Eagle’s Minimum Essential Medium (EMEM; (Seroglob) supplemented with 7% heat-inactivated fetal bovine serum (FBS, Seraglob), 2 mmol/L Glutamax (Gibco), 100 μg/mL Streptomycin and 100 IU/mL Penicillin (Gibco). Whereas the MDCK-I cells were maintained in EMEM, supplemented with 5% heat-inactivated FBS, 100 μg/mL Streptomycin and 100 IU/mL Penicillin (Gibco). Both cell lines were propagated at 37°C in a humidified incubator with 5% CO_2_.

### Viruses

Influenza D virus (D/Bovine/Oklahoma/660/2013) was inoculated on MDBK cells and propagated in infection medium (EMEM, supplemented with 0.5% Bovine Serum Albumin (Sigma-Aldrich), 15 mmol/L of HEPES (Gibco), 100 μg/mL Streptomycin and 100 IU/mL Penicillin (Gibco), and 1 μg/mL Bovine pancreas-isolated acetylated trypsin (Sigma-Aldrich)). Infected MDBK cultures were maintained for 96 hours at 37°C. The ICV strain C/Johannesburg/1/66 was inoculated on MDCK-I cells and propagated in infection medium for 96 hours at 33°C. Virus containing supernatant was cleared from cell debris through centrifugation for 5 minutes at 500x *rcf* before aliquoting and storage at -80°C.

### Human airway epithelial cell culture

Primary human bronchial cells were isolated from patients (>18 years old) undergoing bronchoscopy or pulmonary resection at the Cantonal Hospital in St. Gallen, Switzerland, in accordance with our ethical approval (EKSG 11/044, EKSG 11/103 and KEK-BE 302/2015). Isolation and culturing of primary human bronchial epithelial cells was performed as previously described (39), with the minor modification of supplementing the BEGM with 10 µmol/L Rho associated protein kinase inhibitor (Y-27632, Abcam).

### Viral replication in hAEC cultures

The hAEC cultures were inoculated with 10.000 TCID_50_, or 32 hemagglutination units, of either IDV or ICV. The viruses where incubated for 1.5 hours at temperatures indicated in a humidified incubator with 5% CO_2_. Afterwards, inoculum was removed and the apical surface was washed thrice with Hanks Balanced Salt Solution (HBSS, Gibco), after which the cells were incubated at the indicated temperatures in a humidified incubator with 5% CO_2_. The infection was monitored as previously described, during which progeny virus was collected by incubating the apical surface with 100 μL HBSS 10 minutes prior to the time point. Collected apical washes were stored 1:1 in virus transport medium for later quantification (39).

### Virus titration by tissue culture infectious dosis 50 (TCID50)

MDBK cells were seeded at a concentration of 40.000 cells per well in a 96-cluster well plates. The following day, medium was removed and cells were washed once with PBS and replaced with 50 μL of infection medium. Virus containing samples were ten-fold serial diluted in infection medium, from which 50 μL was added to the MDBK cells in six technical replicates per sample. The inoculated cells were incubated for 72 hours at 37°C in a humidified incubator with 5% CO_2_, where after they were fixed by crystal violet to determine the titre according to the protocol of Spearman-Kärber (40).

### Quantitative real-time reverse transcription PCR

For quantification of the viral kinetics of IDV and ICV, viral RNA was extracted from 50 μL apical wash using the NucleoMag VET (Macherey-Nagel), according to manufacturer guidelines, on a Kingfisher Flex Purification system (Thermofisher). Two microliters of extracted RNA was amplified using TaqMan™ Fast Virus 1-Step Master Mix (Thermofisher) according to the manufacturer’s protocol using the forward primer 5′- AACCTGCTTCTGCTTGCAATCT-3′, reverse 5′-AACAATGAACAGTTACCGCATCA-3′ and probe 5′-FAM-AGACCTGTCTAAAACTATTT-BHQ1-3′ targeting the P42-segment of ICV (AM410042.1). Whereas for the P42-segment of IDV (KF425664.1) the forward 5′- ATGCTGAAACTGTGGAAGAATTTTG-3′, reverse 5′- GGTCTTCCATTTATGATTGTCAACAA-3′ and probe 5′-FAM-AAGGTTTATGTCCATTGTTTCA-BHQ1-3′ were used. A standard curve of the P42-segment of Influenza C or D virus, cloned in pHW2000 plasmid, was included to interpolate the amount of genomic equivalents (41). Measurements and analysis were performed using an ABI7500 instrument and software package (ABI).

### Immunofluorescence of hAEC cultures

The hAEC cultures were formalin-fixed and stained for immunofluorescence as previously described (39). For the detection of IDV-positive cells, hAEC cultures were stained with a custom generated rabbit polyclonal antibody directed against the nucleoprotein (NP) of the prototypic D/Bovine/Oklahoma/660/2013 strain (Genscript). Alexa Fluor^®^ 647-labeled donkey anti-Rabbit IgG (H+L) (Jackson Immunoresearch) was applied as secondary antibody. For the characterization and quantification of the cell tropism, hAEC cultures were stained with the custom generated polyclonal rabbit anti-NP (Genscript), mouse Anti-β-tubulin IV (AB11315, Abcam) and goat anti-ZO1 (AB99642, Abcam). As secondary antibodies were the following antibodies used; Alexa Fluor^®^ 488-labeled donkey anti-mouse IgG (H+L), Cy3-labeled donkey anti-goat IgG (H+L) and Alexa Fluor^®^ 647-labeled donkey anti-Rabbit IgG (H+L) (Jackson Immunoresearch). In the case of ICV, hAEC cultures were stained with human IVIg (Sanquin, the Netherlands), mouse Anti-β-tubulin IV (AB11315, Abcam), rabbit anti-ZO1 (617300, Thermofisher). Using Alexa Fluor^®^ 488-labeled donkey anti-mouse IgG (H+L), Alexa Fluor^®^ 594-labeled donkey anti-human IgG (H+L) and Alexa Fluor^®^ 647-labeled donkey anti-Rabbit IgG (H+L) (Jackson Immunoresearch) as secondary antibodies. All samples were counterstained using 4',6-diamidino-2-phenylindole (DAPI, Thermofischer) to visualize the nuclei. The immunostained inserts were mounted on Colorforst Plus microscopy slides (Thermofischer) in Prolong diamond antifade mountant (Thermo Fischer) and overlaid with 0.17 mm high precision coverslips (Marienfeld). The Z-stack images were acquired on a DeltaVision Elite High-Resolution imaging system (GE Healthcare Life Sciences) using a step size of 0.2 µm with a 60x/1.42 oil objective. Images were deconvolved and cropped using the integrated softWoRx software package and processed using Fiji (ImageJ) and Imaris version 9.1.3 (Bitplane AG, Zurich, Switzerland) software packages.

## Data presentation

Data was plotted using GraphPad Prism 7 and figures were assembled in Adobe Illustrator CS6.

## Acknowledgements

We like to thank Feng Li from the South Dakota University, United States, for providing the IDV (D/Bovine/Oklahoma/660/2013) and Georg Herrler, University of Veterinary Medicine Hannover, Germany for providing ICV (C/Johannesburg/1/66). This study was supported by the Swiss National Science Foundation (project 179260).

## Biographical sketch first author.

Melle Holwerda is a PhD-student at the Department of Infectious diseases and Pathobiology, Vetsuisse faculty at the University of Bern. His focus lays in the characterization of emerging viruses and developing tools to study how these pathogens interact with the host.

